# On adaptation to a mesophilic environment and the chaperone network in Archaea

**DOI:** 10.1101/2025.06.02.657341

**Authors:** Mathieu E. Rebeaud

**Affiliations:** Laboratory of Statistical Biophysics, Institute of Physics, School of Basic Sciences, École Polytechnique Fédérale de Lausanne – EPFL, 1015 Lausanne, Switzerland

**Keywords:** DnaK, Evolution, Temperature, Archaea, Chaperones

## Abstract

A prevailing hypothesis for the emergence of life on Earth holds that it might have originated in hydrothermal vents, where the environmental conditions, although physically and chemically extreme (acidity, lack of oxygen, high pressure, very high temperature), vary very little. According to this view, single-celled organisms appeared under these conditions subsequently began to colonize all aquatic environments, followed by terrestrial ones. Here, I study the proteomes of more than 250 reference proteomes of archaea as well as those of a few non-reference Promethearchaeati (ASGARD), which have an optimal growth temperature of between 10°C and 100°C. I found a correlation between the chaperome present in these organisms, and in particular the presence/absence of the HSP70 family (DnaK-DnaJ-GrpE, KJE for brevity) and the optimal growth temperature. These findings suggest that appearance of HSP70s in mesophilic living conditions was the key to greater adaptability of the organisms, to their ability to colonize different environments and, ultimately, led to the appearance of eukaryotes.

## Introduction

Since life first appeared on Earth at the bottom of the oceans near hydrothermal vents, some 3.5-4 billion years ago, constant evolution has driven organisms to colonize the entire Earth, taking forms as varied as single-celled bacteria and archaea, plants, fungi, sponges, right up to metazoans like cats, blue whales and human [1, 2]. Life appeared in conditions that can be described as extreme if we compare them with the environment in which most eukaryotes tend to evolve. Although conditions near hydrothermal vents, characterized by high acidity, lack of oxygen, extreme pressure, and temperatures up to 120°C, may seem extreme, that thrive under such conditions, they are nonetheless remarkably stable, unchanging, and non-oxidizing due to their location at the bottom of the ocean [2].

It has already been studied in the past that organisms living in environments where the temperature is considered extreme generally have proteins that are optimally adapted to these conditions [3, 4], and therefore have very little chance of further evolving. Singer and Hickey [4] showed that thermophilic organisms living in elevated growth temperatures are imposed with selective constraints at all three molecular levels; the nucleotide content, codon usage and amino acid composition. It has been established that the amino acid composition of thermophilic proteins confers enhanced stability of the folded state at high temperatures [4, 5]. Moreover, several studies have shown that hyperthermophiles such as the model organisms *Thermus thermophilus* [6] and *Sulfolobus acidocaldarius* [7] have lower mutation rates than their mesophilic counterparts. Indeed, neutral mutations can be compensated in mesophilic conditions, and as a consequence mesophilic organisms tend to evolve more rapidly and diversify more easily[8]. Instead, the same mutations likely become deleterious in the extreme physical and chemical conditions of hyperthermophiles. Furthermore, in mesophilic bacteria [9] and archaea [8], this selective pressure appears to be more pronounced for highly expressed genes and proteins[10, 11].

The phylogenomics of 10,575 bacterial and archaeal genomes has recently demonstrated the proximity between the two domains of prokaryotic life, as well as the numerous exchanges between them, mainly through horizontal gene transfer (HGT) [12-14]. The constant presence of HGTs also affects « housekeeping genes », emphasizing the importance of both inter- and intra-domain HGT[15]. Because of the shuffling of the gene pool, tracing the appearance or disappearance of certain genes requires in-depth study of gene families, such as chaperones like HSP60 (GroEL in bacteria, thermosomes in archaea) or the main component of the chaperone network, HSP70 (DnaK for the most conserved in prokaryotes). According to several studies, HSP70s first appeared in mesophilic bacteria, then was transferred by HGT to archaea [1, 16, 17].

DnaK is considered to be the central hub of the chaperone network, and in *Escherichia coli* about 700, mostly cytosolic, proteins usually interact with DnaK under standard growth conditions [18]. Furthermore, a recent study on the bacterium *Myxococcus xanthus* demonstrated that DnaK duplication and specialization in bacteria correlates with increased proteome complexity [19]. DnaK has already been determined to increase mutational robustness, as well as to allow proteins that are its obligate clients to evolve faster than those that are not by supporting their folding [20]. Therefore, the presence of DnaK, and more in general of the Protein Quality Control (PQC) system, has profound long-term effects on genome evolution [21-23]. These studies provide evidence that even a single protein, or a limited set of proteins (the chaperones), can have a disproportionate effect on the evolution of the proteome.

In this study, I wanted to investigate the relationship between the size of the proteome of over 250 mesophilic or thermophilic archaea as a function of their chaperome, of the presence or absence of HSP70 and the associated system (DnaJ, GrpE and also HSP90 like HtpG and HSP100 like ClpB) [24, 25], and of their optimal growth temperature. The aim is to extend our understanding of the relationship between an organism’s complexity, its environment and its chaperone system, this time with a focus on archaea (and some bacteria), unlike our previous study that covered less specifically the whole Tree of Life [1].

## Results

### Selection of archaea and comparison between optimal growth temperature and GC content of 16S RNA of prokaryotes

Around twenty years ago, Kimura and colleagues carried out a selective phylogenetic analysis targeting the 16S rRNA genes of thermophiles and hyperthermophiles, demonstrating the correlation between optimal growth temperature and the GC composition of 16S rRNA [26]. More recently, Hu and colleagues carried out the same type of analysis and concluded that there was a correlation between the GC composition of hyperthermophiles and growth temperature [27]. To investigate this relationship further, I analyzed the 16S rRNA sequences of selected prokaryotes finding, as expected, that that GC content in 16S increases with temperature, likely due to the thermodynamic stability of GC-rich sequences [27, 28] (Fig. 1B). I next looked at the correlation between proteome size and optimal growth temperature, finding that organisms adapted to higher temperatures have on average smaller proteomes (Fig. 1A), possibly due to constraints on protein stability or metabolic efficiency. These trends appear stronger in archaea, which is in keeping with the observation that most hyperthermophiles organisms are of the archaea domain [29] (archaea in general seem to be better adapted to more extreme conditions).

**Figure 1:**
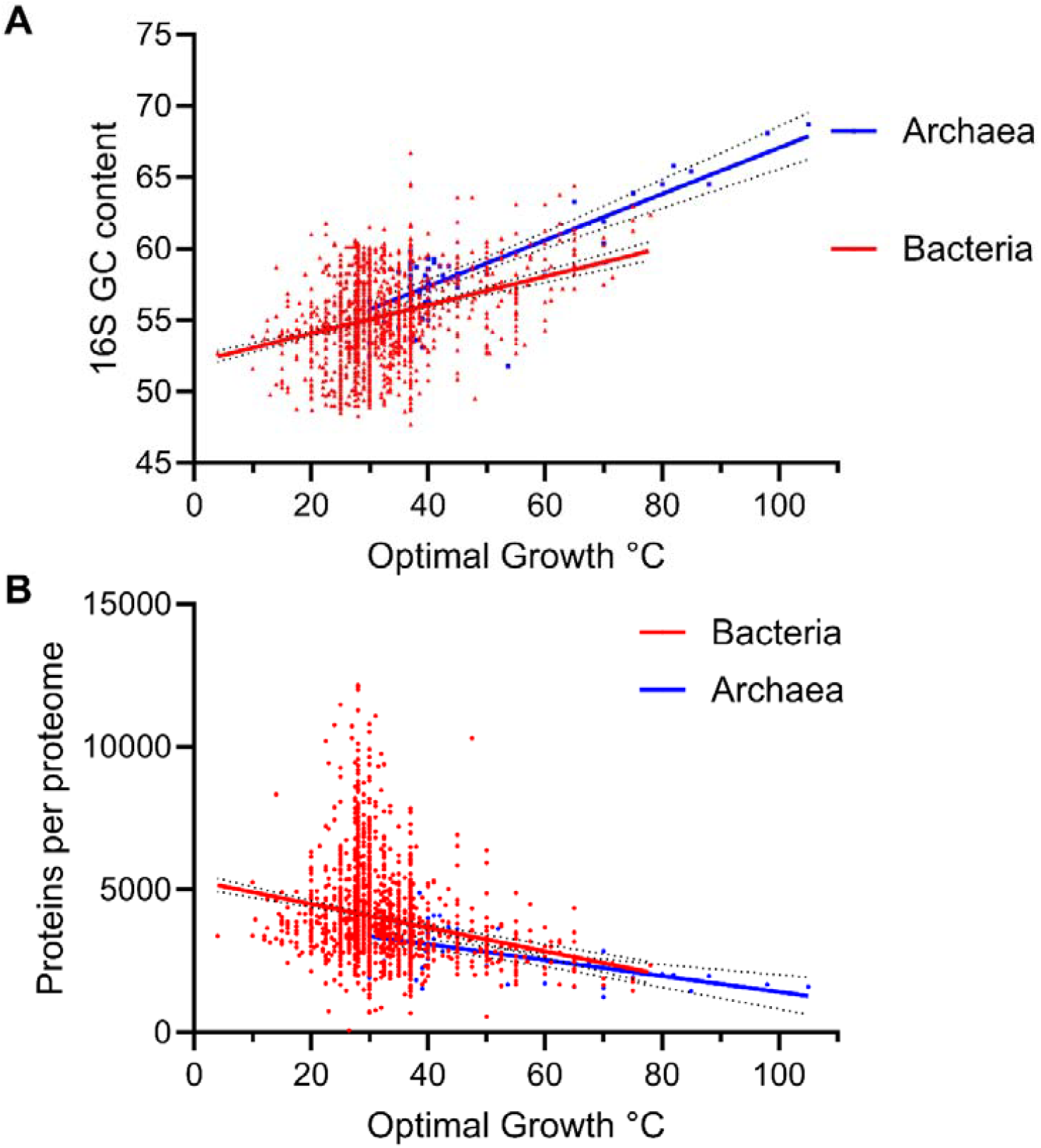
Scatter plot of proteins per proteome and 16S GC content of selected organisms from the Tempura and Uniprot databases. A: Mean values for proteins of archaeal proteomes (blue) are 2920 and for bacterial (red) 4026. Fitted trend lines show a negative correlation between optimal growth temperatures and proteomes size. B: Bacteria and archaea show a positive correlation between GC content and optimal growth temperatures. Archaea (blue) have a steeper trend line than bacteria (red), suggesting a stronger relationship between temperature and GC content in archaeal species. Trend lines are simple linear regression between the x and y axis calculated with PRISM.

While the GC content and the gene count (proteome) provide a bird’s eye view of the genetic fingerprint of adaptation to high temperature, they do not directly address how cells that evolved at very disparate temperatures can ensure protein stability and function. This question invokes the role of the PQC, which is essential for protein folding and stress response. Thus, I next examined the correlation between the presence or absence of specific chaperone families and growth temperatures in archaea.

Hyperthermophilic organisms, and therefore those that are potentially the most interesting for answering the questions posed here, are those from the archaeal domain based on the analysis. The Tempura database which was already curated was used as a reference to see which prokaryotes are the most interesting one, but it contains very few archaeal organisms, so it will be necessary to use another database (BacDive [30]) and select organisms manually from UniProt references proteomes to refine the results for archaea. I thus collected 234 archaeal proteomes (see Methods and Supplementary Information) whose proteome average size was 2763 ± 960, comparable to the first (Sup Fig. 1), which is 2920± 880 (see supplementary information) [30].

**Supplementary Figure 1:**
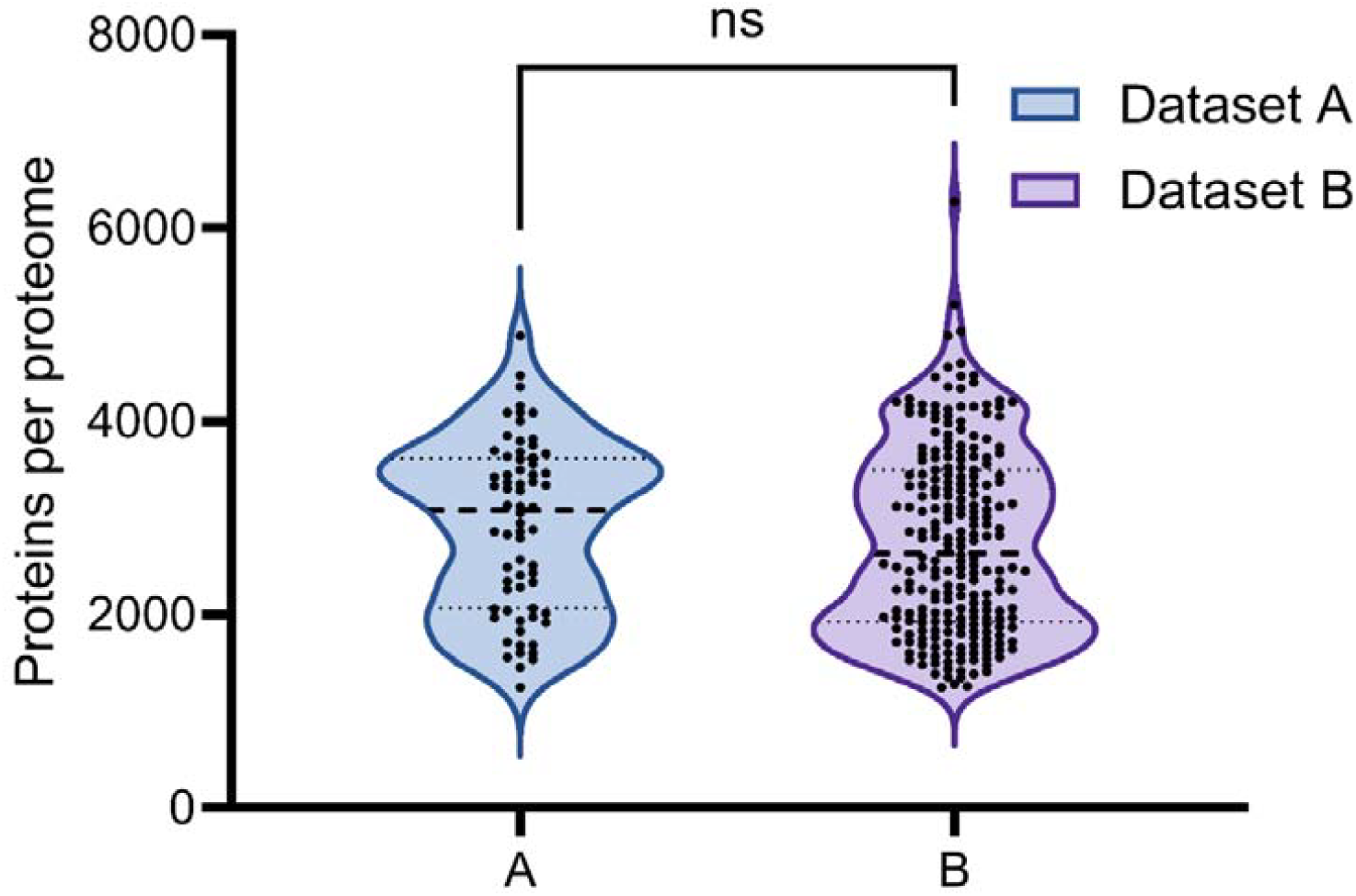
Size comparison of the mean proteomes size between the two datasets used in this study. Mean proteins number in dataset A is 2920 and dataset B 2763. Black dots represent individual values. Black dots representing individual proteomes. Statistical comparisons (t-test) between datasets are denoted by asterisks, ns: p value >0.1.

The correlation between proteome size and optimal growth temperature again suggests that hyperthermophilic archaea have smaller proteomes (Fig. 2), confirming our previous results (Fig.1B).

**Figure 2:**
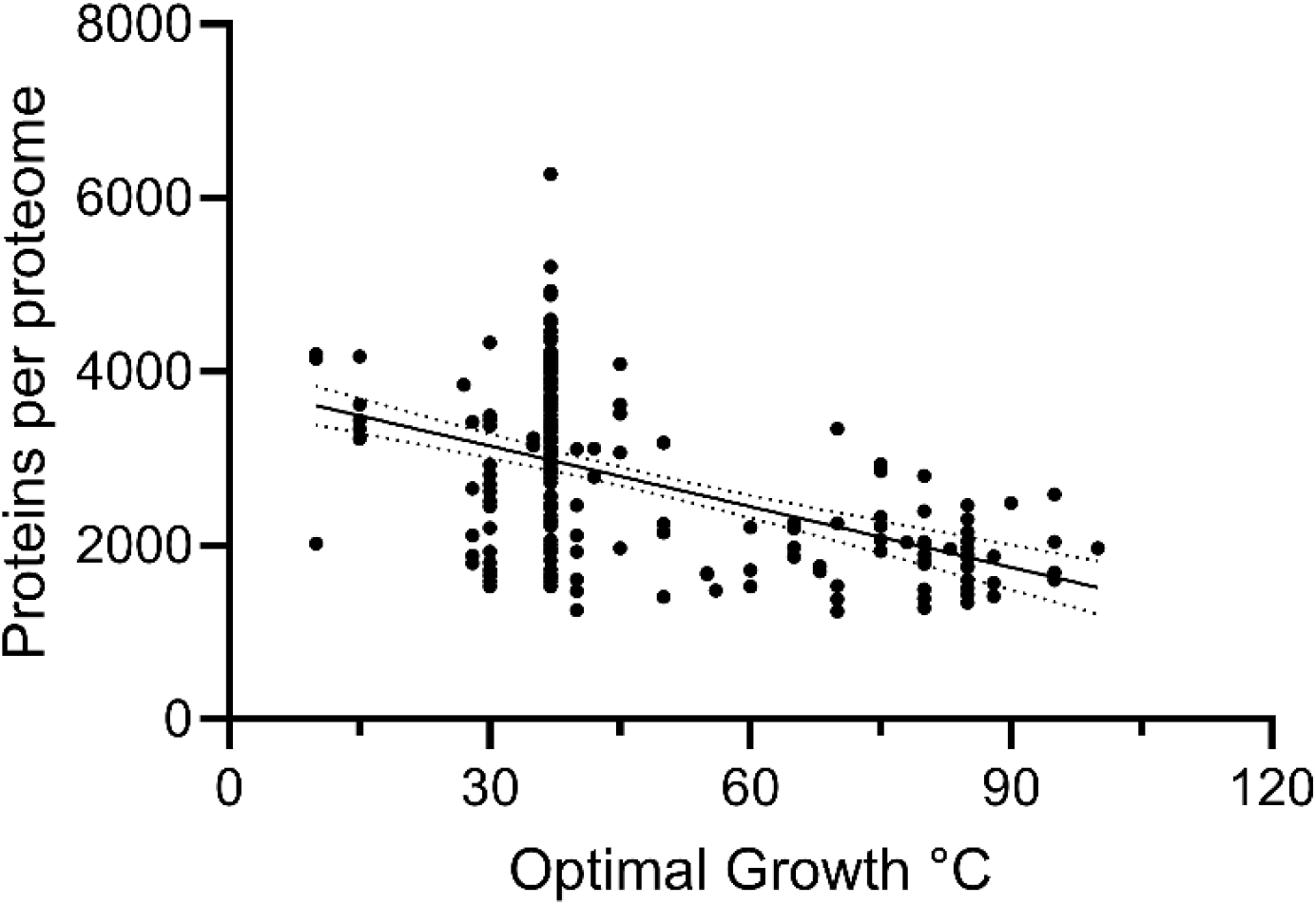
Scatter plot of proteins per proteome for selected archaea. Mean value for proteins of archaeal proteomes is 2763. Fitted trend lines show a negative correlation between optimal growth temperatures and proteomes size. Trend lines are simple linear regression between the x and y axis calculated with PRISM.

This correlation between optimal growth temperature actually tend to recapitulate the taxonomic classification of different archaeal groups, which show distinct trends in proteome size and growth temperature. Some groups, like Halobacteriales, display a broader range of proteome sizes while preferring lower temperatures. Others, such as Thermococci, have smaller proteomes and grow at higher temperatures (Fig. 3).

**Figure 3:**
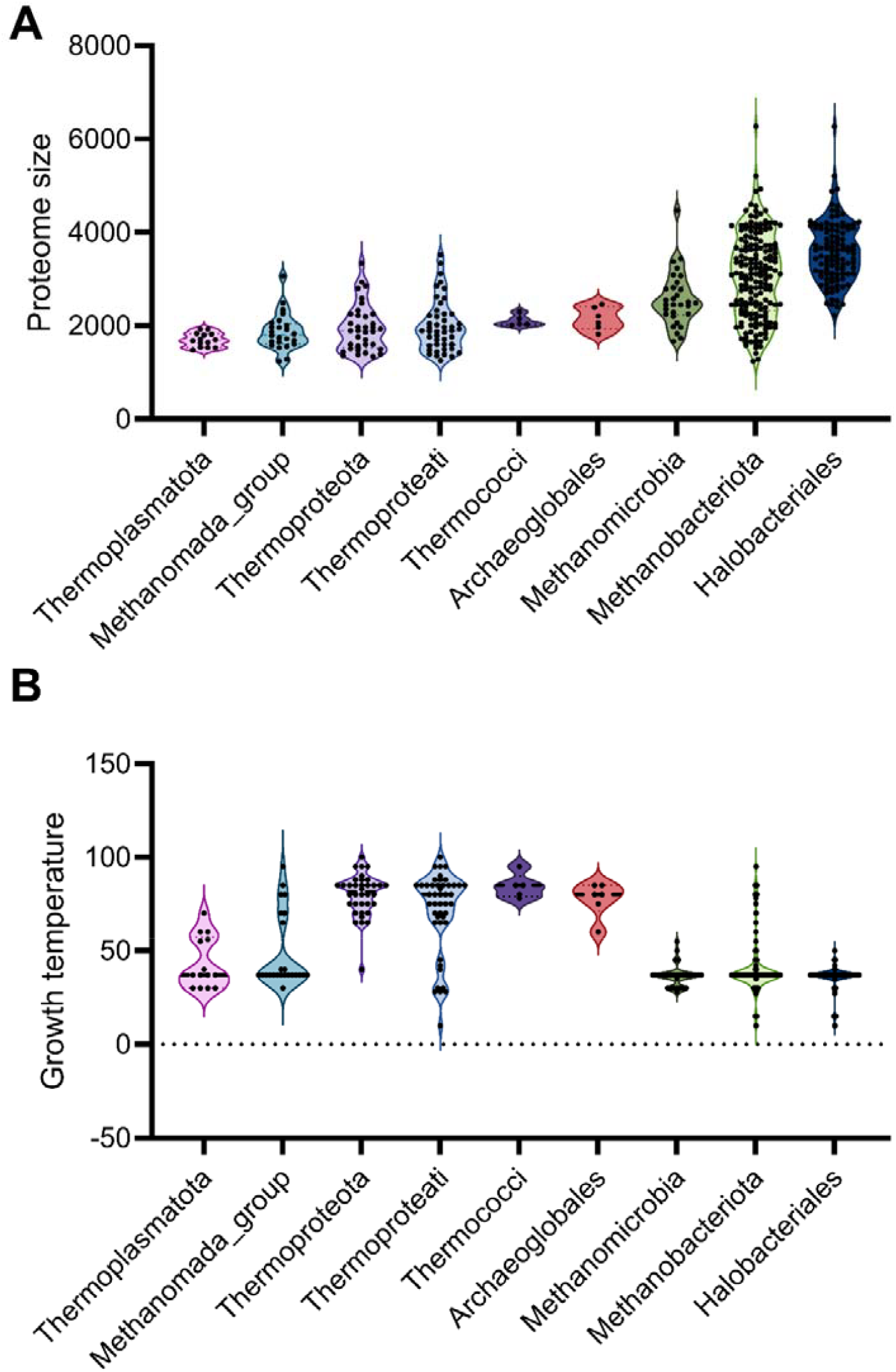
Comparison of different microbial groups based on their mean proteome size (A) and mean growth temperature (B). A: Proteome sizes vary significantly across groups, with some, like Halobacteriales, having larger ranges. B: Differences in growth temperatures, where thermophilic groups (e.g., Thermoprotei, Thermococci) have higher mean growth temperatures than others. Black dots represent individual values.

### Chaperones and selected genes network in archaea

A detailed survey of the presence of the different chaperones and the proteasome subunits (psmA and psmB, since the proteasome is required for cell growth and stress responses [31]) is shown in Fig.4. The vast majority of all the archaea analyzed here (∼98-100%) possess one or more subunits of the thermosomes (chaperonins) [32], sHSPs [33], prefoldins [34] (here called TPS system) and 20S proteasomes [31], attesting for their crucial relevance at all temperatures. As a matter of fact, prefoldins, chaperonins and sHSPs are considered to be among the most conserved in archaea and constitute the oldest chaperone system that, according to several bioinformatics analyses, was already part of the Last Universal Common Ancestor (LUCA) [2]. Using these protein families also enabled us to check the databases and the robustness of the analysis, given that thermosomes are thought to be ubiquitously present and highly expressed [32].

One chaperone system showed instead a striking correlation with optimal growth temperature: the DnaK-DnaJ-GrpE (KJE) system, which is mostly associated with mesophilic species (Fig. 4; Sup Fig. 2). In particular, attesting for the stringent relation between DnaK and DnaJ, they are always present (or absent) together. Instead, GrpE was absent in a few organisms that have DnaJ and DnaK, but was never present in their absence, a feature that has already been observed in the past in several parasitic organisms [35]. Nonetheless I cannot exclude sequencing problems, since the physiological interest in losing GrpE, which greatly enhances the efficiency of the KJ system, would be evolutionarily difficult to explain [1, 35]. It is also notable that the proteomes of archaea that possess DnaK, and the KJE system in general, are larger (Fig. 4 and Fig. 5).

**Figure 4:**
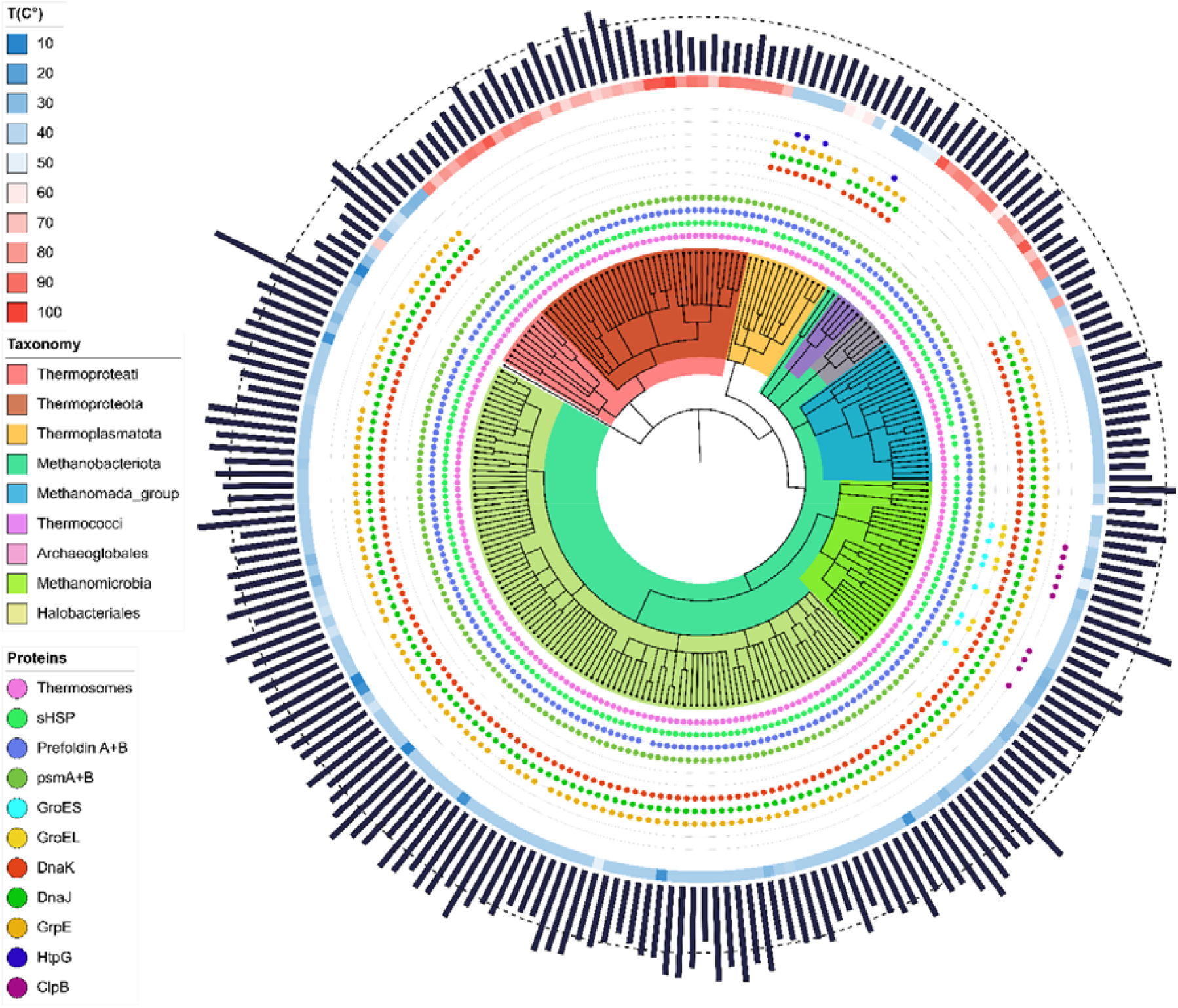
Taxonomic tree of the selected 234 archaeal organisms. Different taxonomy has been represented on the inner tree; Thermoproteati (light red), Thermoproteota (maroon), Thermoplasmatota (orange), Methanobacteriota (sea green), Methanomada group (sea blue), Thermoccoci (purple), Archaeoglobales (pink, grey), Methanomicrobia (lime) and Halobacteriales (pear). On the outside of the tree are different classes of chaperones and proteases represented by the presence (colored circle) or absence of the protein; Thermosomes (pink), sHSP (sea green), Prefoldin A/B (blue), Proteasome subunit A/B (green), GroES (light blue), GroEL (yellow), DnaK (red), DnaJ (green), GrpE (yellow), HtpG (deep blue) and ClpB (purple). Then an heatmap of the optimal growth temperature of the organisms between 10 (blue) to 100 °C (red). On the outer side are proteome size of each organism (black bars) with the mean size (dotted line). This tree has been constructed using the NCBI taxonomy [31] of selected organisms and visualized in iTOL [32].

**Figure 5:**
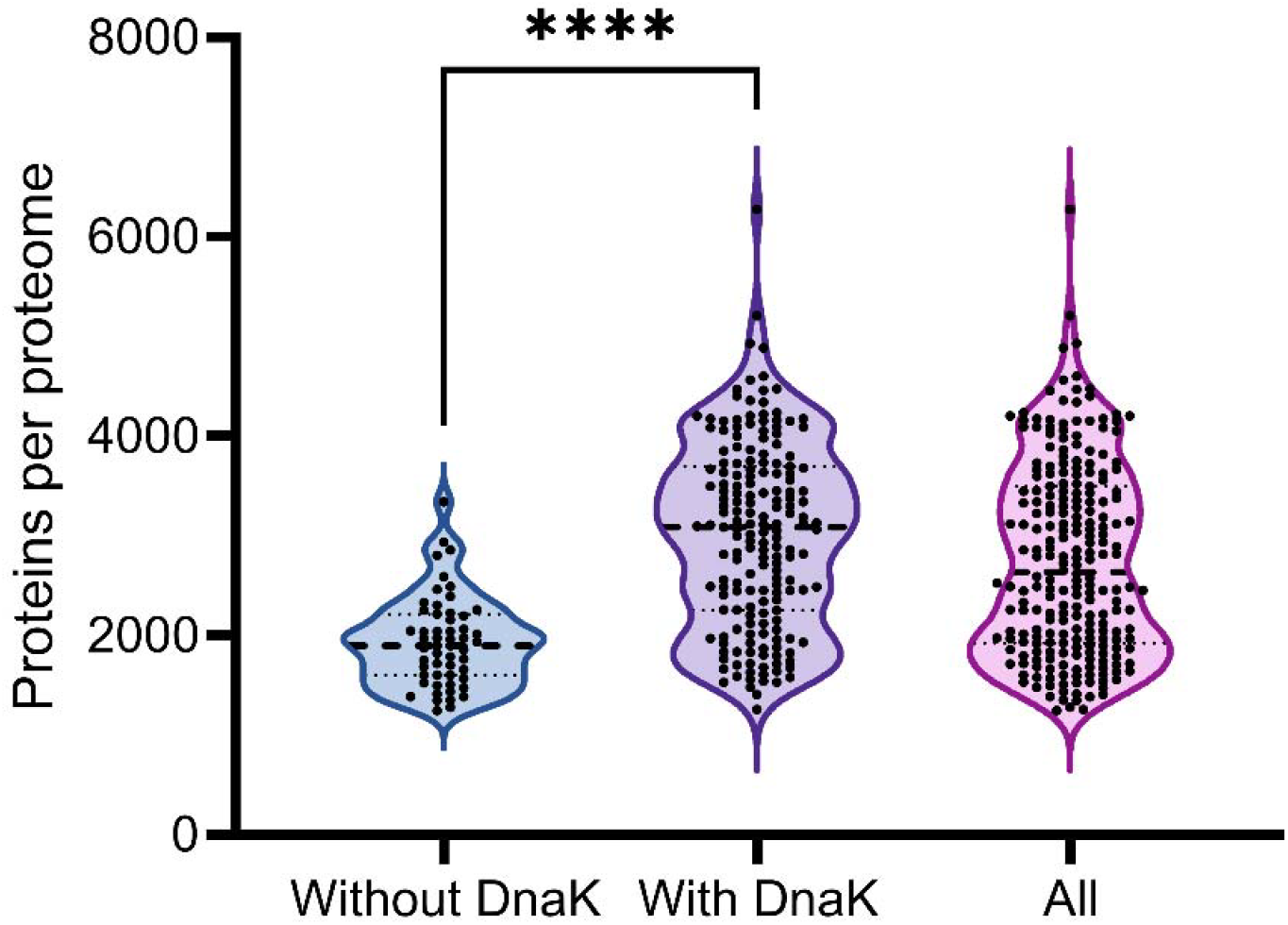
Mean archaeal proteome size depending on the presence or absence of DnaK. Mean archaeal proteome size of all archaea in the subset is 2763 proteins. Without DnaK, 1936 and with DnaK 3023. The y-axis represents the number of proteins per proteome, ranging from 0 to 8000. Black dots representing individual proteomes. Statistical comparisons (t-test) between datasets are denoted by asterisks, ***: p value >0.0001.

The Methanomicrobia class, to which *Methanosarcina acetivorans* [36] (lime clade in Fig. 4 and supplementary information) belongs, is a class of archaea, some of whose members possess group I and II chaperonins, namely the thermosomes and GroELs [37, 38]. Since archaea have acquired many bacterial genes through HGT, it’s not surprising that an archaeon of this class, such as *Methanosarcina mazei* [39], possesses both classes of chaperonins, the latter having different substrate specificity. In addition to the group I chaperonins, this specific group of archaea seems to have acquired the AAA+ chaperone of the HSP100 family (ClpB) for several of its members (purple dots on Fig. 4, supplementary information). This suggests that members of this mesophilic group have undergone a fairly large number of HGTs for the chaperone network. At least one member of this group has been closely studied for the presence of ClpB within its proteome [40]. A final point that raises questions is the presence of ClpB only in Methanomicrobia, which have also acquired GroELS. Is it the specificity of the environmental niche [41] in which these archaea live that has enabled successive HGTs with bacteria that are also present? Studying the prokaryotic communities that cohabit in this type of niche could provide an answer, as well as studying what other genes might have been transferred. Another finding that deserves to be explored in more detail is the presence in some Thermoplasmatota organisms of the chaperone HSP90 (HtpG, deep blue dots on Fig. 4), which is generally accepted to be absent from most archaea [42].

**Supplementary Figure 2:**
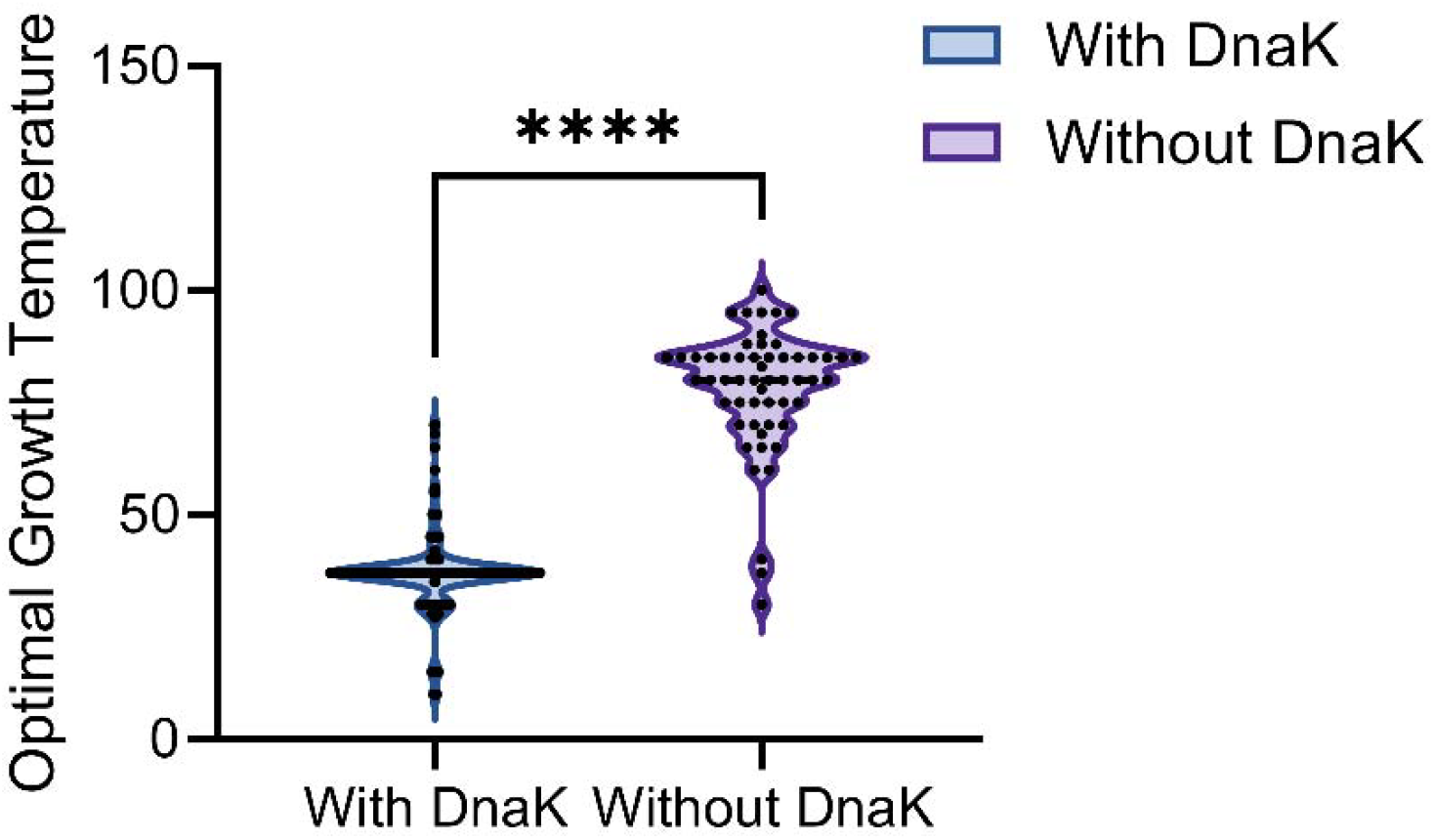
Optimal growth temperature of organisms with or without DnaK. Mean for organisms with DnaK is 36.75°C and for those without 78.18 °C. Black dots representing individual proteomes. Statistical comparisons (one-way ANOVA) between datasets are denoted by asterisks, ***: p value <0.0001.

In recent years, several new members of the archaea domain have been discovered, among which the ASGARD family [43, 44] which were renamed and classified in the Promethearchaeati kingdom [45] by Imachi and colleagues recently. This group is currently thought to be the closest to the archaea that fused with bacteria during endosymbiosis to become the Last Eukaryotic Common Ancestor, LECA [46]. Although most of the available proteomes of these new archaea are not considered reference proteomes, and thus were not considered in the analysis above, I selected some of the most complete and annotated ones in order to characterize their chaperome, optimal growth temperatures, and the relative size of their proteomes. While most of these taxa have not been validly described based on their nomenclatural status [47], *Prometheoarchaeum syntrophicum* MK-D1 being the only reference organism. I reprise here the same analysis presented earlier, but on this more specific dataset, showing that, relative to other archaea, Promethearchaeati tend to possess larger proteomes (Sup Fig. 3), consistent with most of them supposedly being mesophilic or moderate thermophilic according to recent studies [48-50].

**Supplementary Figure 3:**
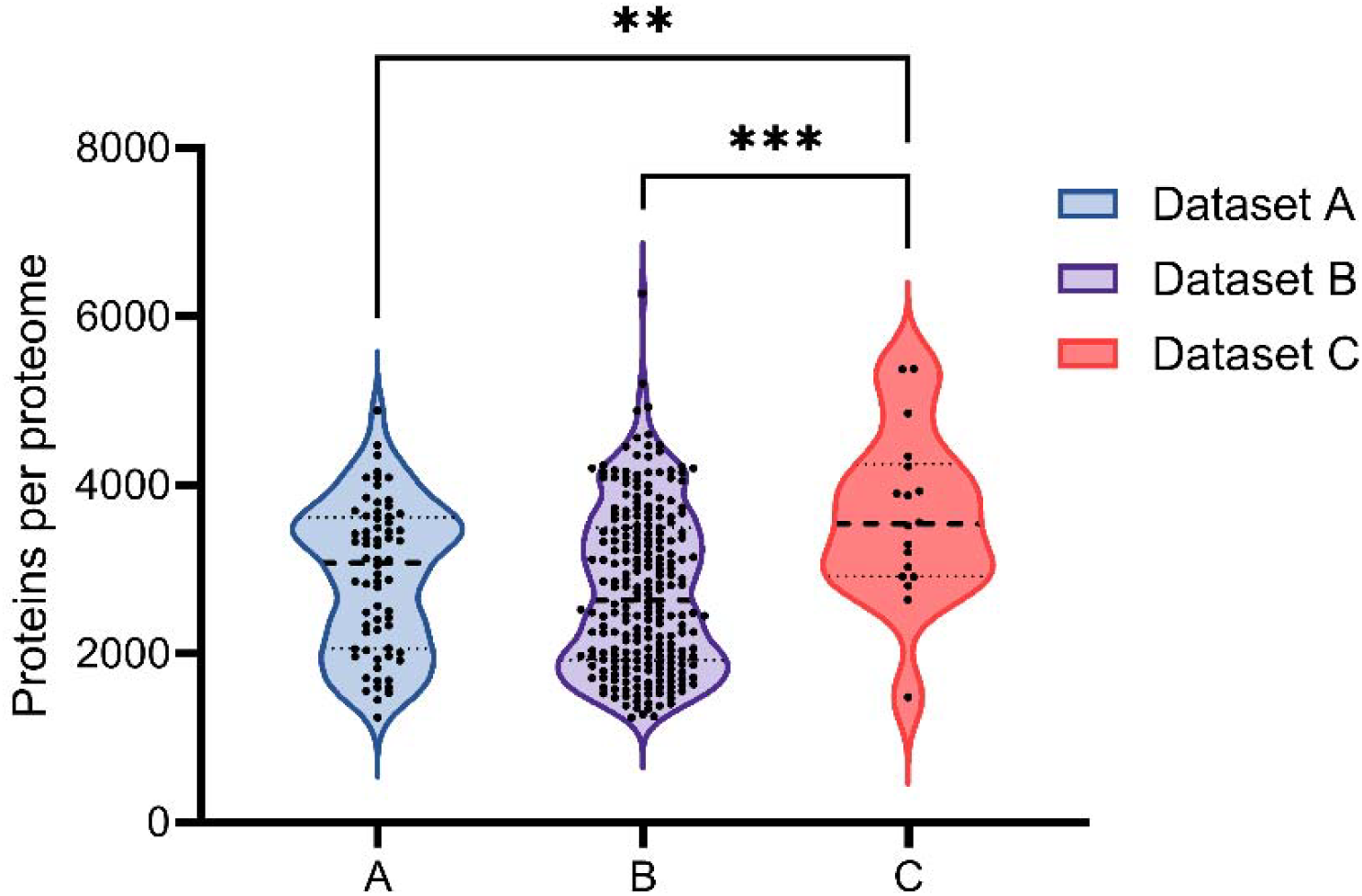
Size comparison of the mean proteomes size between the three datasets used in this study. Mean proteins number in dataset A is 2920 (±880), dataset B 2763 (±960) and dataset C 3626 (±988). The y-axis represents the number of proteins per proteome, ranging from 0 to 8000. Black dots representing individual proteomes. Statistical comparisons (one-way ANOVA) between datasets are denoted by asterisks, **: p value <0.01, ***: p value <0.001.

I then correlated this analysis with the chaperone composition, finding that they all possess the KJE system (Fig. 6), and some have HSP90 (HtpG) or HSP100 (ClpB). That these mesophilic organisms, considered to be the closest relatives of eukaryotes [51], have a more robust chaperone system is not surprising, as proteome complexity and eukaryotic-related complexity found in Promethearchaeati [52] often go hand in hand with chaperone system extension [1]. Interestingly, it appears that HGT from bacteria did not occur for GroEL/S chaperonins in the archaea selected here (Fig. 6), but due to the small set of genomes analyzed here, I refrain from drawing general statements.

**Figure 6:**
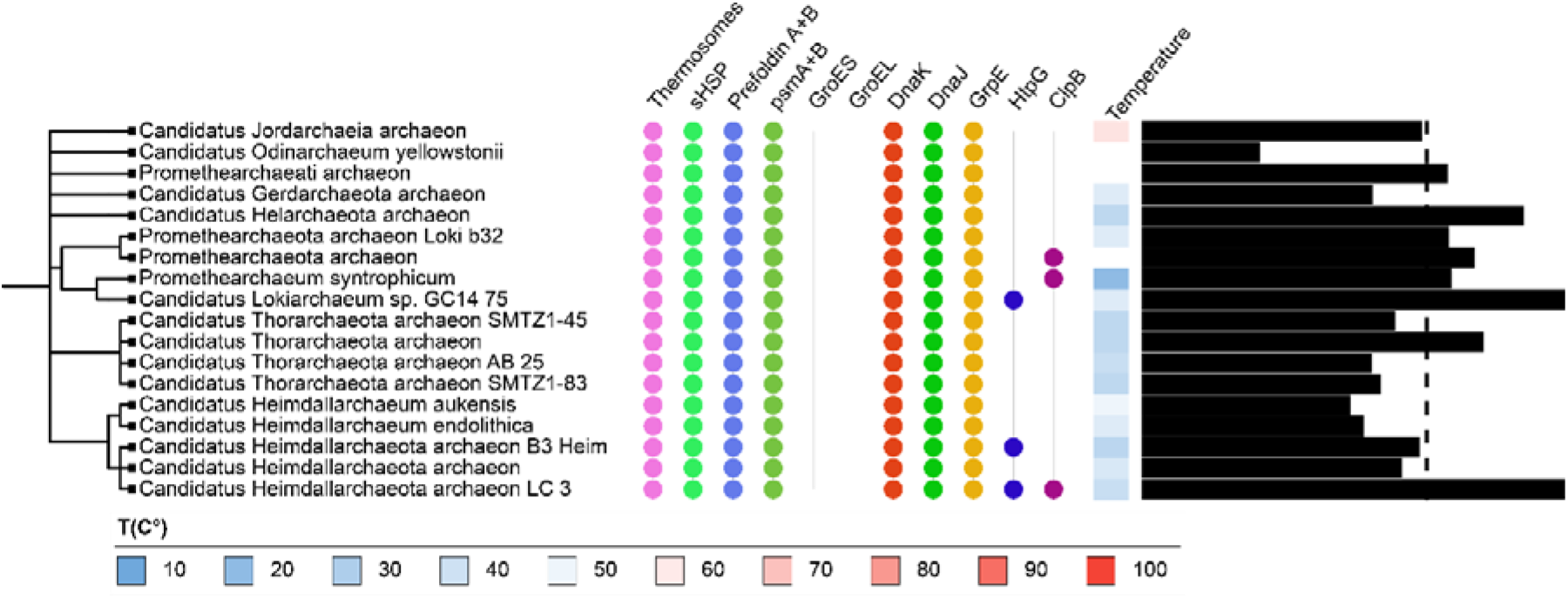
Taxonomic tree of the selected 18 Promethearchaeati organisms. On the outside of the tree are different classes of chaperones and proteases represented by the presence (colored circle) or absence of the protein; Thermosomes (pink), sHSP (sea green), Prefoldin A/B (blue), Proteasome subunit A/B (green), GroES (light blue), GroEL (yellow), DnaK (red), DnaJ (green), GrpE (yellow), HtpG (deep blue) and ClpB (purple). Then an heatmap of the optimal growth temperature of the organisms between 10 (blue) to 100 °C (red). On the outer side are proteome size of each organism (black bars) with the mean size (dotted line, 3626). This tree has been constructed using the NCBI taxonomy [31] of selected organisms and visualized in iTOL [32].

I’ve then selected one HSP70 sequence from each organism displaying one or more Hsp70 genes (*i*.*e*. 178 archaea from the first dataset and the 18 Promethearchaeati) and they were aligned to verify the sequence homology (see supplementary information). The presence of one or more copies of DnaK or HSP70 in mesophilic or light thermophilic organisms, as well as in all Promethearchaeati, is an interesting marker for the joint hypothesis of the maintenance of proteins that possess otherwise deleterious mutations, as well as proteome expansion [19].

### Cold Shock and RNA chaperones

Adaptation to higher temperatures may influence the acquisition of cold-shock proteins (Csps, like CspA, [53] and supplementary information) in the same manner as the adaptation to cold environment of the bacterium Shewanella [54], where “cold” in this case is defined in relation to the optimum temperature for growth. Likewise, RNA chaperones might also play a crucial role in controlling the structure of RNA molecules in the cell.

Here I’ve found only 114 organisms out of 234 that have CspA-like proteins, most of them being Halobacteriales (all the 106 having at least one copy of these proteins) and all of them being mesophilic. All organisms that have CspA-like proteins have also the DnaK-DnaJ-GrpE system, but the opposite is not true. Interestingly, RNA chaperones apparently function in the same way. A study based on three DEAD box RNA helicases [55] (CsdA, SrmB, RhlB) and the cold shock protein CspA (CspA was also investigated in this study because of its RNA-binding properties (see supplementary information)) might suggest that RNA chaperones and binding proteins might play a fundamental role in RNA evolution [56]. One study analysed a DEAD box RNA helicase homolog in the hyperthermophilic archaea *Thermococcus kodakaraensis* [57] and showed that it was transcribed more dominantly at 60°C than at 85°C and 93°C, indeed hinting that “cold” is relative and that, because of this, cold adaptation in hyperthermophiles might reveal general principles of cold shock response. It would be interesting in the future to compile an inventory of DEAD-boxes and associates in archaea, as has been done in bacteria [58].

### Translation optimization and tRNA Adaptation Index of chaperones and ribosomal proteins

Beyond the presence or absence of specific chaperones, another factor influencing the efficiency of protein folding is translation optimization [59]. In a cell, virtually every amino acid can be encoded by several synonymous codons which are selected for their speed of translation and precision [60, 61]. Codons that are translationally optimized can be recognized by tRNAs more easily and therefore translated more quickly [61]. Since the optimized codons are often found in the most aggregation-prone parts of proteins and in domains or subdomains that are highly conserved evolutionarily [62, 63], the selection of optimal codons to improve translation fidelity must be an important determinant of coding sequence evolution [64]. On top of that, codon usage bias can affect protein stability and adaptation to environmental conditions [65]. I used the method first described by dos Reis [61] to calculate translation efficiency, which takes into account the intracellular concentration of tRNA molecules and the efficiency of each codon-anticodon pair. This method is called the tRNA Adaptation Index (tAI) and has been used extensively to answer many questions relating to the regulation of gene expression and molecular evolution, among others [66-68]. It has since been improved by using organism-specific tRNAs to accurately calculate codon usage for each gene in an organism, making the comparison species-specific with an index called stAI [69], which I used to determine whether a selection of archaeal chaperones are optimized for efficient translation. Using the GtRNAdb [70] to fetch the tRNAs of selected organisms and the genomes available on NCBI [71], I selected a subset of 14 organisms from the 234 archaea with optimal growth between 10 and 100°C. To better compare differences between species, a Z-score was applied to all calculated stAIs (Sup Fig. 4), as there is a disparity in stAIs between and among kingdoms [72], and previous studies have failed to find rules governing the evolution of codon usage mechanism [73, 74]. This allowed me to check whether most organisms have ribosomal and chaperone proteins that have better translation efficiency (or in some cases lower) than the average, which is normally the case for these types of proteins [32, 35] and since codon usage regulates co-translational protein folding [74].

**Supplementary Figure 4:**
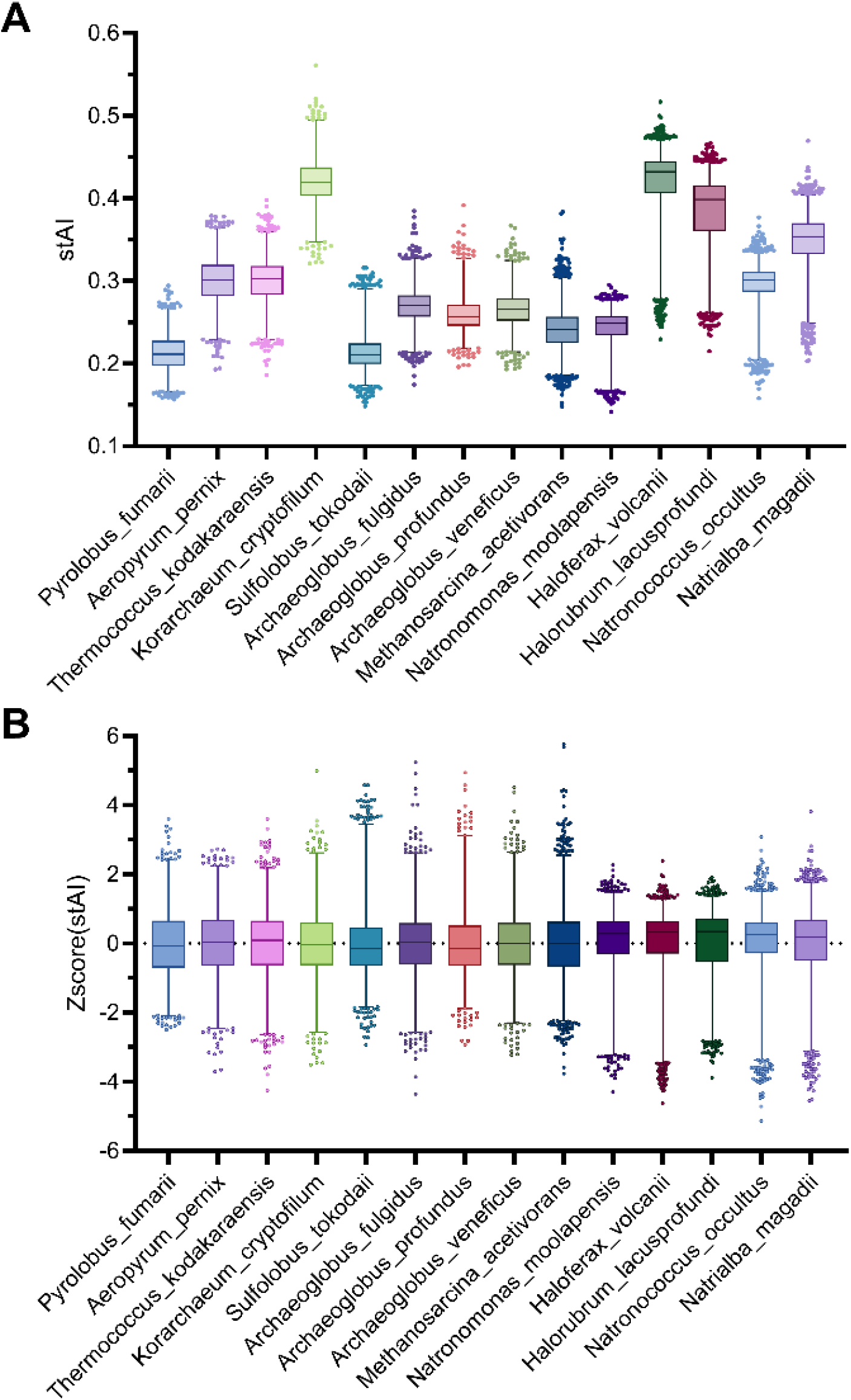
Specie specific tRNA Adaptation Index. A: specie tRNA Adaptation Index of 14 selected organisms based on the tRNA of the GtRNAdb and the complete genome on NCBI database. Each gene is given a tRNA Adaptation Index between 0 to 1 based on its translational adaptation. B: same as (A) but with a Z-score applied on each organism so the mean value of each is 0.

To assess whether the patterns of translation optimization observed in archaeal proteomes extended to key protein groups, I specifically evaluated the stAI scores of ribosomal proteins, which are generally among the most highly expressed and efficiently translated proteins in all domains of life. Virtually all the organisms selected have on average ribosomal proteins with better translation efficiency, although there is some discrepancy between proteins within the same organism (Sup Fig. 5), with just two mesophilic and psychrophilic (optimal growth of 10° and 30°, supplementary information), terrestrial archaea (*Natronococcus occultus* and *Halorubrum lacusprofundi*) showing no statistical difference between the whole genome and ribosomal proteins profile. The same analysis has been performed on the model organism *Saccharomyces cerevisiae* for a comparison between a single-celled eukaryote and archaea (Sup Fig. 6). It is interesting to note, as a quality check, that the subunits of the CCTs in yeast all have an almost equivalent level of translational adaptation (Sup Fig. 6).

**Supplementary Figure 5:**
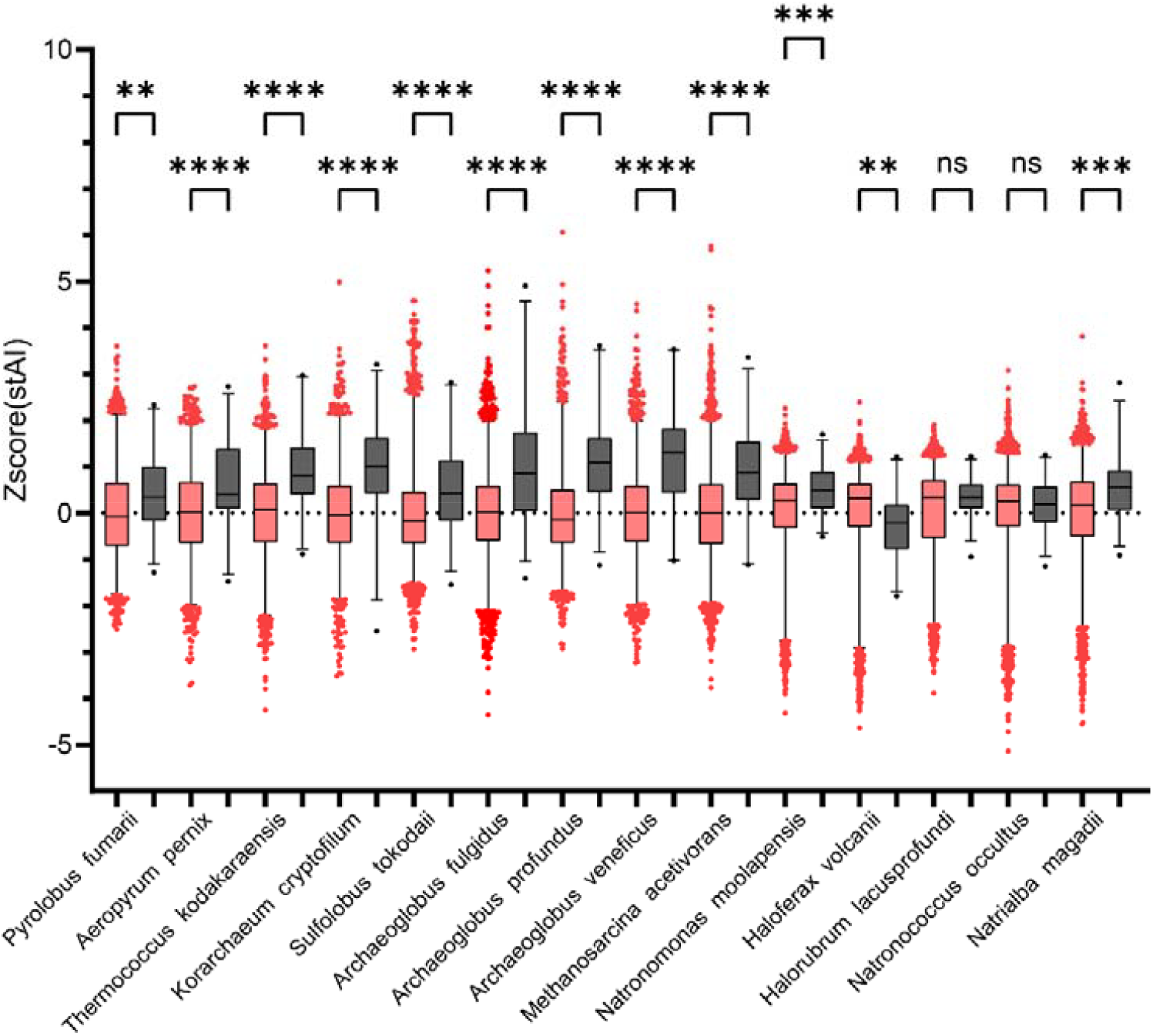
Specie specific tRNA Adaptation Index of selected archaea and Ribosomal proteins. The Z score of specific tRNA Adaptation Index of 14 selected organisms based on the tRNA of the GtRNAdb and the complete genome (red boxes) on NCBI database with comparison to the ribosomal proteins (black boxes) of each of the selected organisms.

**Supplementary Figure 6:**
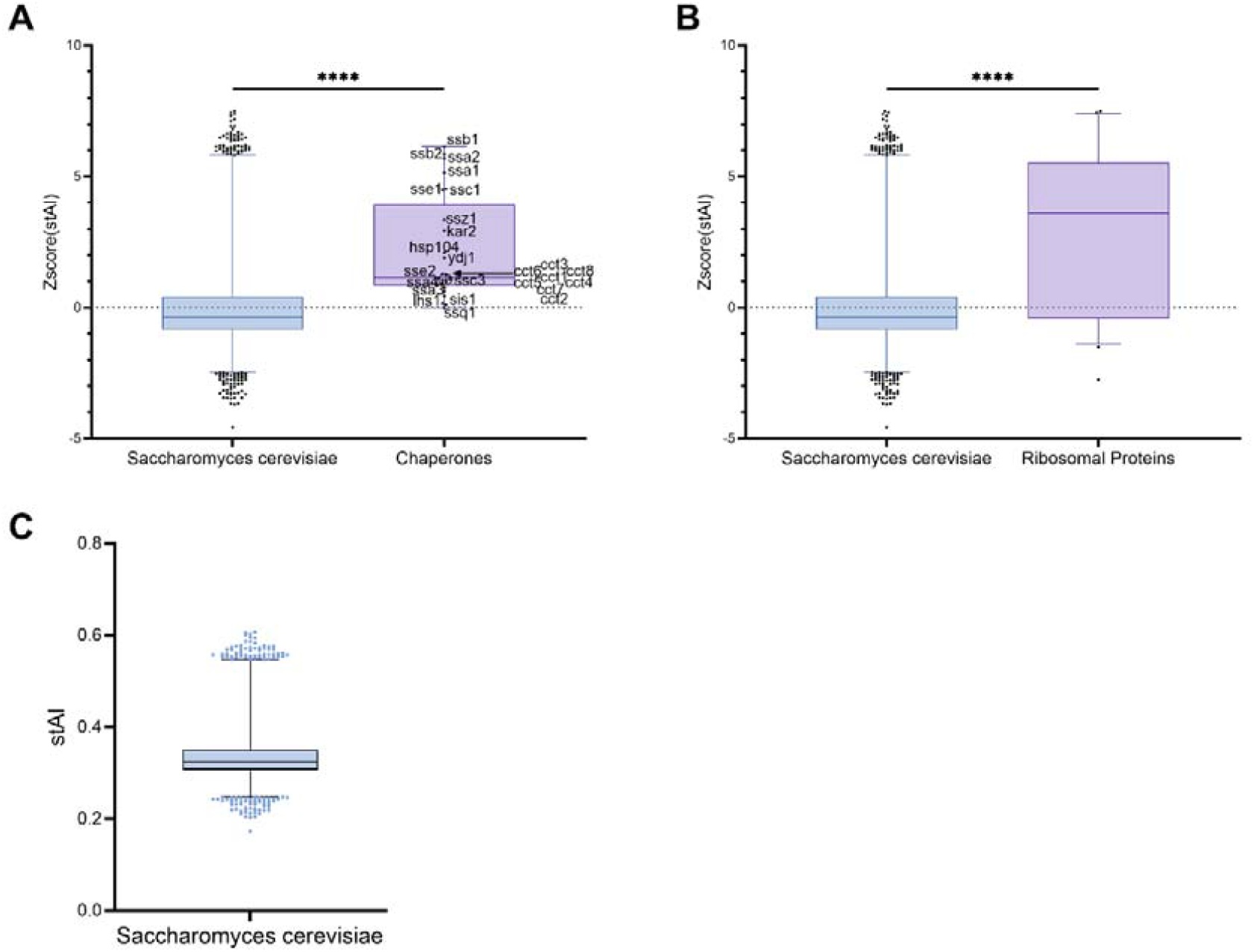
Specie specific tRNA Adaptation Index of *S*.*cerevisiae*. A: The Z-score applied on the stRNA Adaptation Index of *Saccharomyces cerevisiae* organisms based on the tRNA of the GtRNAdb and the complete genome on NCBI database with comparison to HSP70-HSP110 and some selected chaperones of yeast. B: same as A but with the ribosomal proteins instead of chaperones. C: stRNA Adaptation Index of *Saccharomyces cerevisiae* organisms based on the tRNA of the GtRNAdb and the complete genome on NCBI database. Each gene is given a tRNA Adaptation Index between 0 to 1 based on its translational adaptation.

By examining the use of codons by species as an alternative means of expression level, it seems that each organism has adapted some of their chaperones according to the families and subunits available (Fig. 7 and Sup Fig. 5). The majority of hyperthermophilic or thermophilic organisms depend solely on the TPS system (thermosomes, prefoldins and small HSP/HSP20) and most of them exhibit the highest level of translational adaptation in these archaea. For example, in *Pyrolobus fumarii*, which grows at 100°C, the two thermosomes subunits, as well as a sHSP and the beta subunit of Prefoldin, are the most highly adapted. Interestingly, it seems that sHSPs are very often those with the greatest disparity between the most and least adapted when we look specifically at the chaperones selected here. Most of the mesophilic and psychrophilic archaea (like *Halorubrum lacusprofundi* and *Natronococcus occultus*) that possess the KJE system also rely on the TPS system, which are also often better adapted. Organisms with DnaK rely on its presence if I look at the level of translational adaptation of this protein, but the same cannot be said for their respective proteostasis systems, as can be deduced from the analysis of gene sets and codon usage patterns, which has been already seen in bacteria [16, 35]. Although most chaperones have a higher-than-average level of translational adaptation, it remains mostly relatively modest in the selected archaea compared with yeast (Sup Fig. 6).

**Figure 7:**
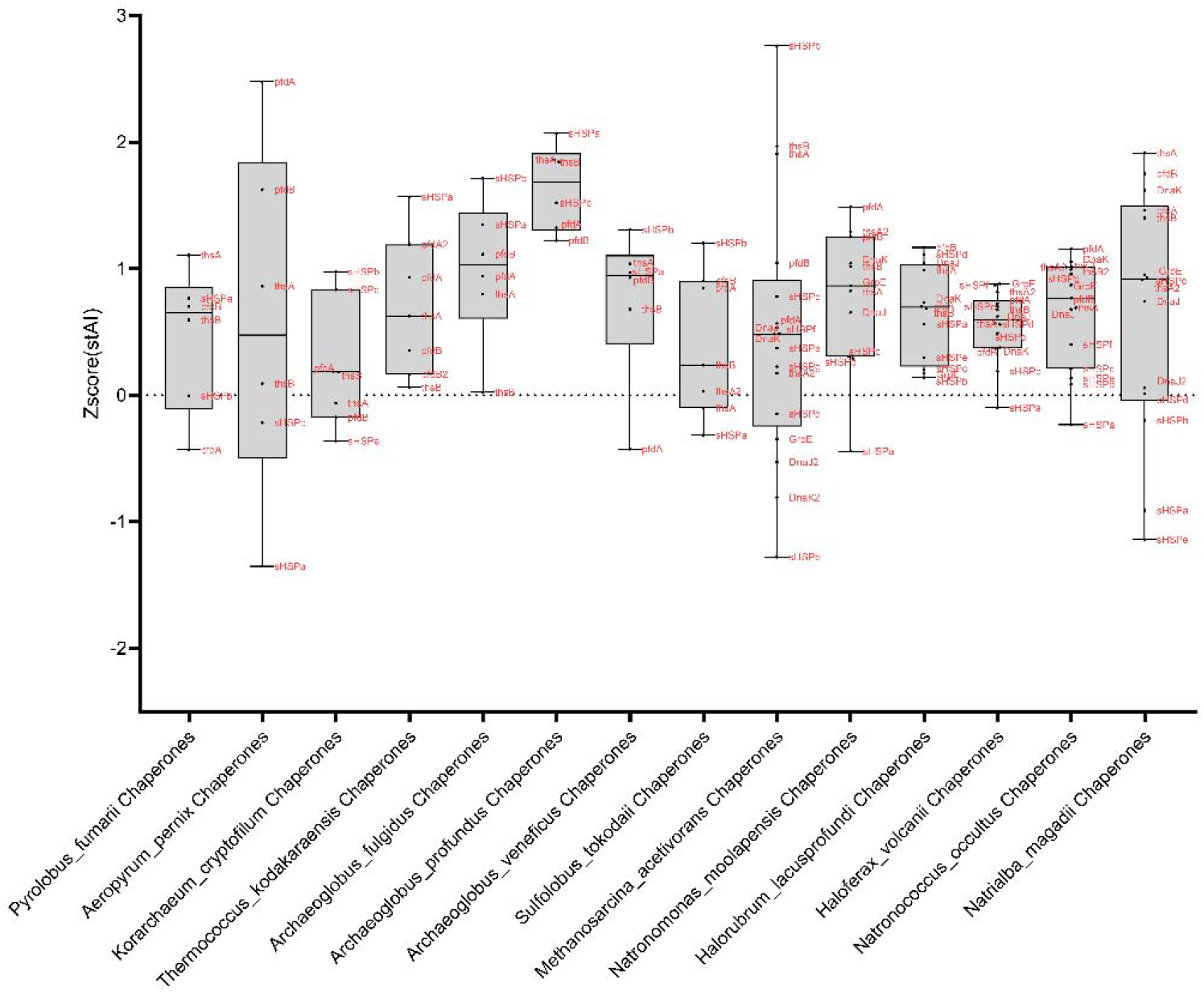
Chaperones tRNA Adaptation Index of selected archaea. Selected chaperones (Dnak-DnaJ-GrpE for the KJE system, Thermosomes subunits (tshA, B), Prefoldins (pfdA, B) subunits and sHSps for the TPS system) are plotted (red names) for each organism.

Analysis of codon usage adaptation reveals that archaeal species appear to have adapted their chaperone systems according to the specific families and subunits available to them (Fig. 7 and Sup. Fig. 5). Notably, the majority of hyperthermophilic and thermophilic archaea rely primarily on the TPS system, comprising thermosomes, prefoldins, and small heat shock proteins (sHSPs/HSP20). These chaperones often show the highest levels of translational adaptation within these organisms. For instance, in *Pyrolobus fumarii*, which thrives at 100°C, both thermosome subunits, one sHSP, and the β-subunit of prefoldin exhibit the strongest adaptation signals.

Interestingly, among the chaperones analyzed, sHSPs frequently display the greatest disparity in translational adaptation levels, with only some variants being highly optimized. In contrast, while many mesophilic and psychrophilic archaea, such as *Halorubrum lacusprofundi* and *Natronococcus occultus* possess the KJE system, they also retain components of the TPS system, which, in these cases, tend to show higher levels of adaptation.

Organisms encoding DnaK exhibit strong translational adaptation for this specific chaperone, suggesting functional dependence. However, this pattern is not consistently observed across the entirety of their proteostasis networks, as inferred from codon usage and gene set analyses, an observation that mirrors findings in bacterial systems [16, 35]. While most chaperones in the selected archaeal species exhibit above-average levels of translational adaptation, their optimization generally remains modest when compared to model eukaryotes such as yeast (Sup. Fig. 6).

## Discussion

It has been proposed that the DnaK chaperone system (KJE) plays a crucial role in Protein Quality Control for adaptation to a mesophilic environment [16, 75]. Most of the mutations happening without any chaperone control are mostly deleterious or neutral, and very few are positive, but the KJE (and HtpG) and GroEL/S systems can actively participate to mutational robustness [3, 22, 76-78]. Indeed, some studies have proposed that by turning deleterious mutations into neutral ones, thus not purged by purifying selection [79], chaperones might increase the evolutionary capacity of their clients, with respect to the one of non-chaperone clients [78, 80]. As a consequence, the ability of clients to withstand mutations enables them to gain functional importance in the cell and ultimately perform a more important role [20, 81]. Likewise, one could speculate that proteins with important functions that have received several mutations need to be supported by chaperones for their correct folding, or risk being purged. It is noteworthy that one family of thermophilic and hyperthermophilic bacteria, the Aquificaceae, possess the KJE system[82, 83], a specificity that is sufficiently surprising that previous studies have noted the absence - or presence - of the system in several members of this family[35].

Another type of adaptation to a mesophilic environment involves genes playing a role in the regulation of membrane fluidity and membrane composition [84, 85] and transport across membranes, as a large proportion of HGT from bacteria belong to two of these metabolic categories [75, 86-89].

Finally, based on the Specie specific tRNA Adaptation Index (stAI) results obtained here, most chaperones (or at least one subunit of complexes) have a higher level of translational adaptation than the average protein in each organism, regardless of the average stAI level (Fig. 7 and Sup Fig. 5). Nevertheless, in comparison with yeast (Sup Fig. 6), it seems that archaeal chaperones generally have a lower level of translational adaptation, even regarding DnaK, than their distant cousins, the eukaryotic HSP70s in yeast (Ssa 1-4, Ssb, Ssc).

Overall, these results highlight the essential role of chaperones in the formation of archaeal proteomes and their adaptation to extreme environmental conditions. In particular, the strong correlation between the KJE system and mesophilic lifestyles suggests that its acquisition may have been a key factor in enabling archaea to thrive outside extreme temperature zones. These results contribute to a better understanding of how molecular chaperones influence evolutionary trajectories.

Ultimately, the conundrum that emerges from this and previous studies is a bit of a chicken-and-egg paradox: is it because of the presence of DnaK that organisms could evolve outside extreme temperature zones, or is it the presence of less extreme temperature that enabled them to acquire DnaK and then evolve more rapidly?

## Conclusion

In summary, I have analyzed over 250 archaeal proteomes to make the widest possible comparison with quality proteomes that are as complete as possible, so as to limit as far as possible the absence of proteins due to poor sequencing or lack of completion. Most of the evidence I have found here points in the general direction that the absence of the KJE system correlates with reduced proteome size and growth temperature in the selected organisms. Furthermore, the absence of KJE combined with the growth medium also allows a hypothesis on the combination of the two parameters that clear organisms of proteins that acquire mutations too rapidly due to the absence of KJE [20] and the purifying selection [79]. Furthermore, it has been demonstrated that temperature is a major evolutionary factor in archaea, and that highly expressed proteins share similarities with thermophilic proteins [5, 8]. This study therefore extends our knowledge of the relationship between proteome size, optimal growth temperature and the chaperone network in archaea.

## Material and Methods

### Prokaryotic database

The Tempura [90] and BacDive [30] databases were used to retrieve information on optimal growth temperatures for the various organisms, as well as information on the GC content of 16S ribosomal RNA. For missing information and values, publications on the various organisms were retrieved to complete the database. In the case of Promethearchaeati, several papers have been used to infer temperatures of most of them [48, 49].

### Selection of organisms and local datasets creation

The archaea selected to create the local database were retrieved on UNIPROT [91] from the reference proteomes and filtered against the BUSCO [92] Score with less than 10% of the proteome missing. For the first dataset, around 2000 archaeal and bacterial organisms were selected based on the Tempura values. Then, 234 archaea were selected to create the second dataset, to which several organisms from the Promethearchaeati (ASGARD) [45] clade were added in a separate third dataset. The Promethearchaeati proteomes are not reference proteomes, but are those of the archaea closest to the common ancestor of all eukaryotes, LUCA [46]. The FASTA files from each of these organisms have been fetched and assembled with BLAST+ [93] to be used [94] on a local database against selected sequences.

### Establishing presence or absence of proteins in the selected datasets

The two datasets containing exclusively archaea (the 234 archaea and the 18 Promethearchaeati) were then blasted against a selection of proteins from bacteria (Escherichia coli K12, UP000000625), archaea (*Thermoplasma acidophilum*, UP000001024; *Methanothermobacter thermautotrophicus*, UP000005223; *Thermoplasma acidophilum*, UP000001024 and *Methanocaldococcus jannaschii* UP000000805). Each selected protein from each organism was blasted against the two local databases with an E_value of 1e-5 for most of the proteins (1e-2 for sHSP) and then manually curated. Based on the result of the BLAST searches and if the presence of hits in each organism were not present, another BLAST was performed with another closely related protein and/or with a lowered E_value up to 1. Subsequent results were analyzed by hand and the presence or absence of each protein was reported. Finally, the results were compared with the InterPro [95] database to check that the proteins obtained, and the domains present were correctly annotated. All proteins used can be found in the Supplementary information. To ensure that some of the proteins were correct, an alignment was carried out using MUSCLE/MAFFT [96, 97] to check the MSA.

### Species-specific tRNA adaptation index (stAI) calculation and Codon Usage Analysis

The tAI measures the extent to which a gene uses codons that correspond to more abundant or more efficiently decoded tRNAs. Here, the stAIcalc [69] technique, a tRNA adaptation index calculator based on species-specific weights, was used via the R Cubar package (<u>10.32614/CRAN.package.cubar</u>). More specifically, the different tRNAs were searched for the selected organisms on the GtRNAdb database [70] to obtain a specific weight per organism according to codon usage, then the total genome was downloaded in FASTA format for the nucleotides of each protein via NCBI [71, 98]. The stAI index calculation was then deduced for each organism in total (all proteins) as well as specifically for each protein of interest. A Z score (Zscore of stAI) was also calculated to standardize the adaptation index between organisms.

### Figure creation

The figures were all drawn using Graphpad PRISM version 10 and iTOL [99] for the phylogenetic and/or taxonomic trees.

## Acknowledgement

I would like to acknowledge Pierre Goloubinoff for many stimulating discussions that enabled me to work on several very interesting topics, such as this one. Paolo De Los Rios for reading the manuscript, amending it, thoughtful comments and suggestions on how to improve it and letting me work on this. Lisa Gennai for reading the manuscript and giving me some feedback, and all my colleagues in the LBS lab.

